# Rare genetic variation in Fibronectin 1 (*FN1*) protects against *APOEe4* in Alzheimer’s disease

**DOI:** 10.1101/2024.01.02.573895

**Authors:** Prabesh Bhattarai, Tamil Iniyan Gunasekaran, Dolly Reyes-Dumeyer, Dörthe Jülich, Hüseyin Tayran, Elanur Yilmaz, Delaney Flaherty, Rafael Lantigua, Martin Medrano, Diones Rivera, Patricia Recio, Nilüfer Ertekin-Taner, Andrew F Teich, Dennis W Dickson, Scott Holley, Richard Mayeux, Caghan Kizil, Badri N Vardarajan

## Abstract

The risk of developing Alzheimer’s disease (AD) significantly increases in individuals carrying the *APOEε4* allele. Elderly cognitively healthy individuals with *APOEε4* also exist, suggesting the presence of cellular mechanisms that counteract the pathological effects of *APOEε4*; however, these mechanisms are unknown. We hypothesized that *APOEε4* carriers without dementia might carry genetic variations that could protect them from developing *APOEε4-*mediated AD pathology. To test this, we leveraged whole genome sequencing (WGS) data in National Institute on Aging Alzheimer’s Disease Family Based Study (NIA-AD FBS), Washington Heights/Inwood Columbia Aging Project (WHICAP), and Estudio Familiar de Influencia Genetica en Alzheimer (EFIGA) cohorts and identified potentially protective variants segregating exclusively among unaffected *APOEε4* carriers. In homozygous unaffected carriers above 70 years old, we identified 510 rare coding variants. Pathway analysis of the genes harboring these variants showed significant enrichment in extracellular matrix (ECM)-related processes, suggesting protective effects of functional modifications in ECM proteins. We prioritized two genes that were highly represented in the ECM-related gene ontology terms, *(FN1)* and collagen type VI alpha 2 chain (*COL6A2*) and are known to be expressed at the blood-brain barrier (BBB), for postmortem validation and *in vivo* functional studies. The FN1 and COL6A2 protein levels were increased at the BBB in *APOEε4* carriers with AD. Brain expression of cognitively unaffected homozygous *APOEε4* carriers had significantly lower FN1 deposition and less reactive gliosis compared to homozygous *APOEε4* carriers with AD, suggesting that FN1 might be a downstream driver of *APOEε4*-mediated AD-related pathology and cognitive decline. To validate our findings, we used zebrafish models with loss-of-function (LOF) mutations in *fn1b* – the ortholog for human *FN1*. We found that fibronectin LOF reduced gliosis, enhanced gliovascular remodeling and potentiated the microglial response, suggesting that pathological accumulation of FN1 could impair toxic protein clearance, which is ameliorated with *FN1* LOF. Our study suggests vascular deposition of FN1 is related to the pathogenicity of *APOEε4*, LOF variants in FN1 may reduce *APOEε4*-related AD risk, providing novel clues to potential therapeutic interventions targeting the ECM to mitigate AD risk.

## Introduction

Alzheimer’s disease (AD) is typically characterized clinically with progressive memory impairment and decline in other cognitive domains; however, there is a long pre-symptomatic period without clinical manifestations^1^. At death, pathological hallmarks in brain include extracellular β-amyloid protein in diffuse and neuritic plaques and neurofibrillary tangles made of hyper-phosphorylated tau protein. AD, a progressive neurodegenerative disorder, is currently unpreventable, and, with available drugs only marginally affecting disease severity and progression, remains effectively untreatable. A critical barrier to lessening the impact of late-onset AD (LOAD) is the slow development of drugs that prevent or treat AD due, in part, to an incomplete characterization of the basic pathologic mechanisms. Determining which genes and gene networks contribute to AD could reveal the biological pathways for drug development, and inform the development of genetic testing methods for identifying those at greatest risk for AD. Presence of the *APOEε4* allele is among the most prominent genetic risk factors for AD in white, non-Hispanic populations ^2^, but the associated risks observed in African-Americans and Hispanics are somewhat lower ^3^. Relative risk of AD associated with a single copy of *APOEε4* is 2.5-fold in Caucasians compared to 1.0 and 1.1 in African Americans and Hispanics respectively ^3^. However, in every population, homozygosity for the *APOEε4* allele is associated with increased risk and nearly complete penetrance ^4–6^. *APOE*, a critical player in lipid metabolism and transport, has been extensively studied for its role in Alzheimer’s disease (AD) and other neurodegenerative disorders ^7–9^. The *APOEε4* allele is a well-established risk factor for late-onset AD, with carriers of this allele exhibiting an increased susceptibility to cognitive decline and dementia and earlier age at onset of clinical symptoms. However, within the population of *APOEε4* carriers, there is variability in age of onset and severity of AD symptoms. Some “resilient” or “cognitively normal, unaffected” individuals who carry the *ε4* allele do not develop AD or experience a delayed onset of symptoms. Several potential factors might contribute to the variability in AD risk and presentation among *APOEε4* carriers. Genetic modifier mutations outside of the *APOE* gene might interact with *APOEε4* to influence the risk of AD. *APOEε4* carriers might also be influenced by other risk factors for AD, such as vascular health, inflammation, and metabolic conditions. Interactions between *APOEε4* and these factors could modify the course of the disease. Certain rare protective variants in other genes could offset the risk posed by *APOEε4*.

Amid the well-documented association between *APOEε4* and AD risk, a growing body of evidence suggests intriguing nuances in the effects of this allele, particularly in certain subsets of individuals who defy the expected trajectory of cognitive decline and remain remarkably resilient to neurodegenerative diseases. Notably, heterozygosity of *APOEε4* has incomplete penetrance ^10^, and the polygenic risk of the rest of genome could stratify *APOEε4* carriers into high and low risk strata. In this study we aimed to identify putative protective mechanisms, influenced by genetic modifiers that might counteract the detrimental effects of the *APOEε4* allele. We sought to identify “protective” genetic factors that can modify or reduce the effect of *APOEε4* on AD risk and to identify new pathogenic mechanisms, proteins and pathways that inform development of therapeutic targets and diagnostics.

## Results

### Whole Genome Sequencing identifies putative protective variants in cognitively unaffected elderly APOEε4 carriers

We accessed whole genome-sequencing data in 3,578 individuals from over 700 non-Hispanic White and Caribbean Hispanic families multiply affected by AD (**Table 1**). After harmonization and QC of the WGS data we identified rare (MAF<1% in GnomAD) coding variants in the healthy elderly (over the age of 70) *APOEε4* homozygous carriers that were absent in non-carriers (**Figure 1**). We further prioritized exon coding variants in healthy *APOEε4* carriers that bear the potential to be damaging to the resulting protein product. **Supplementary Tables 1-3** provide lists of candidate variants that were identified in cognitively unaffected elderly *APOEε4* carriers. Our strategy and analysis pipeline are summarized in **Figure 2**. We found 510 variants in 476 genes that were present in at least 1% of *APOEε4* unaffected homozygous carriers (388 in EFIGA/WHICAP and 130 in NIA-FBS and 8 variants found in both datasets) (**Supplementary Tables 1 and 2**). Two mutations (rs116558455 and rs140926439) in the *FN1* gene (Fibronectin-1) were found in healthy elderly *ε4* homozygous carriers in EFIGA/WHICAP and NIA AD-FBS cohorts with MAF=1.85% and 3.33%, respectively (**Table 2**). In Hispanics, rs116558455 was absent in all *APOEε4* carriers with AD. In non-Hispanic whites rs140926439 was absent in homozygous *APOEε4* AD patients but found in 1% of heterozygous patients. Pathway analysis of the genes harboring variants segregating in *APOEε4* carriers identified several biological pathways and molecular functions such “actin binding”, “microtubule binding”, and “extracellular matrix structural constituent” (**Figure 3**). These results suggested a strong correlation with cellular morphologies and the architectural organization of those cells.

**Figure 1:**
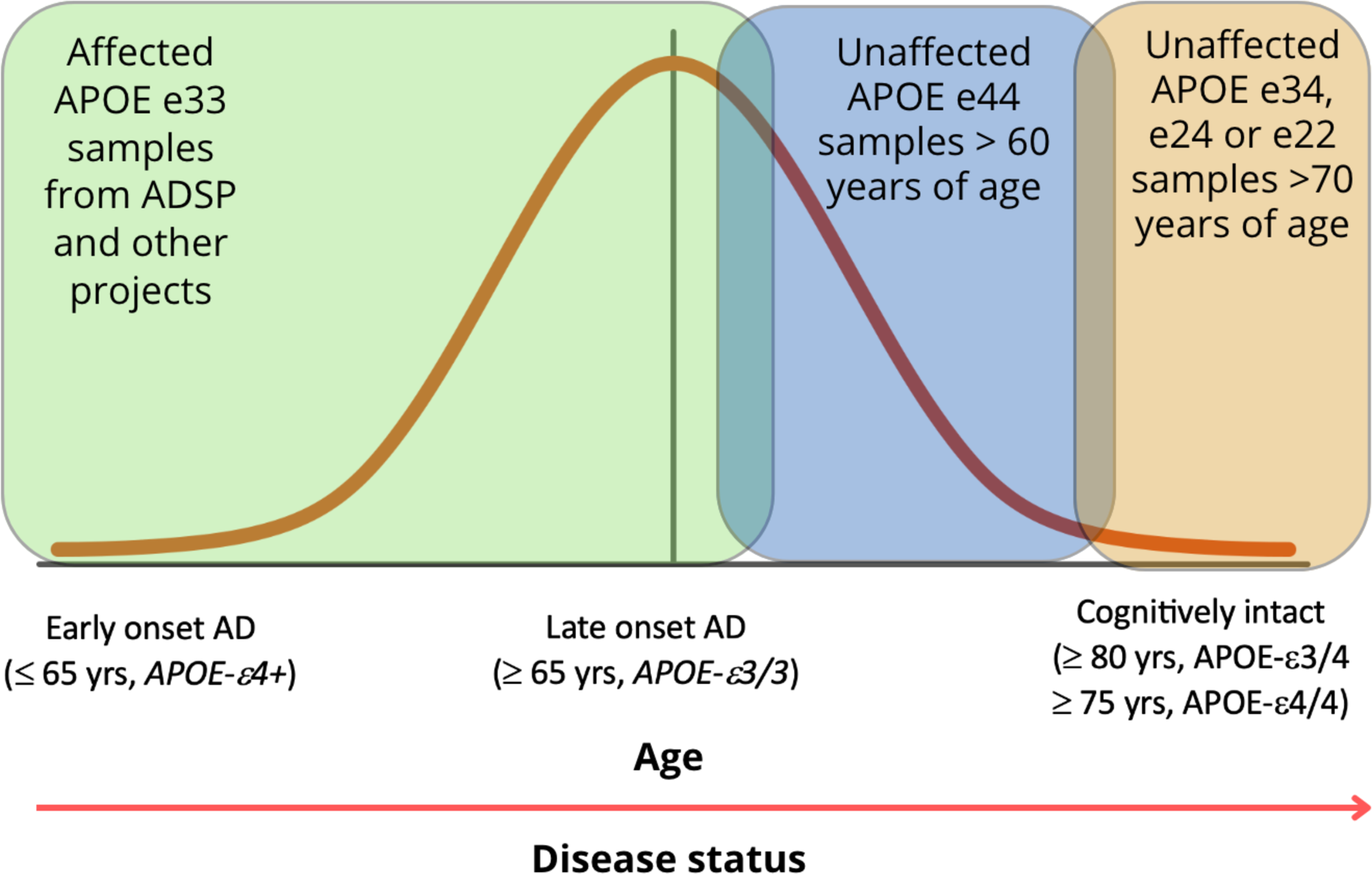
Study Design. Comparison of the genomes of elderly *APOEε4* carriers with non-carriers

**Figure 2:**
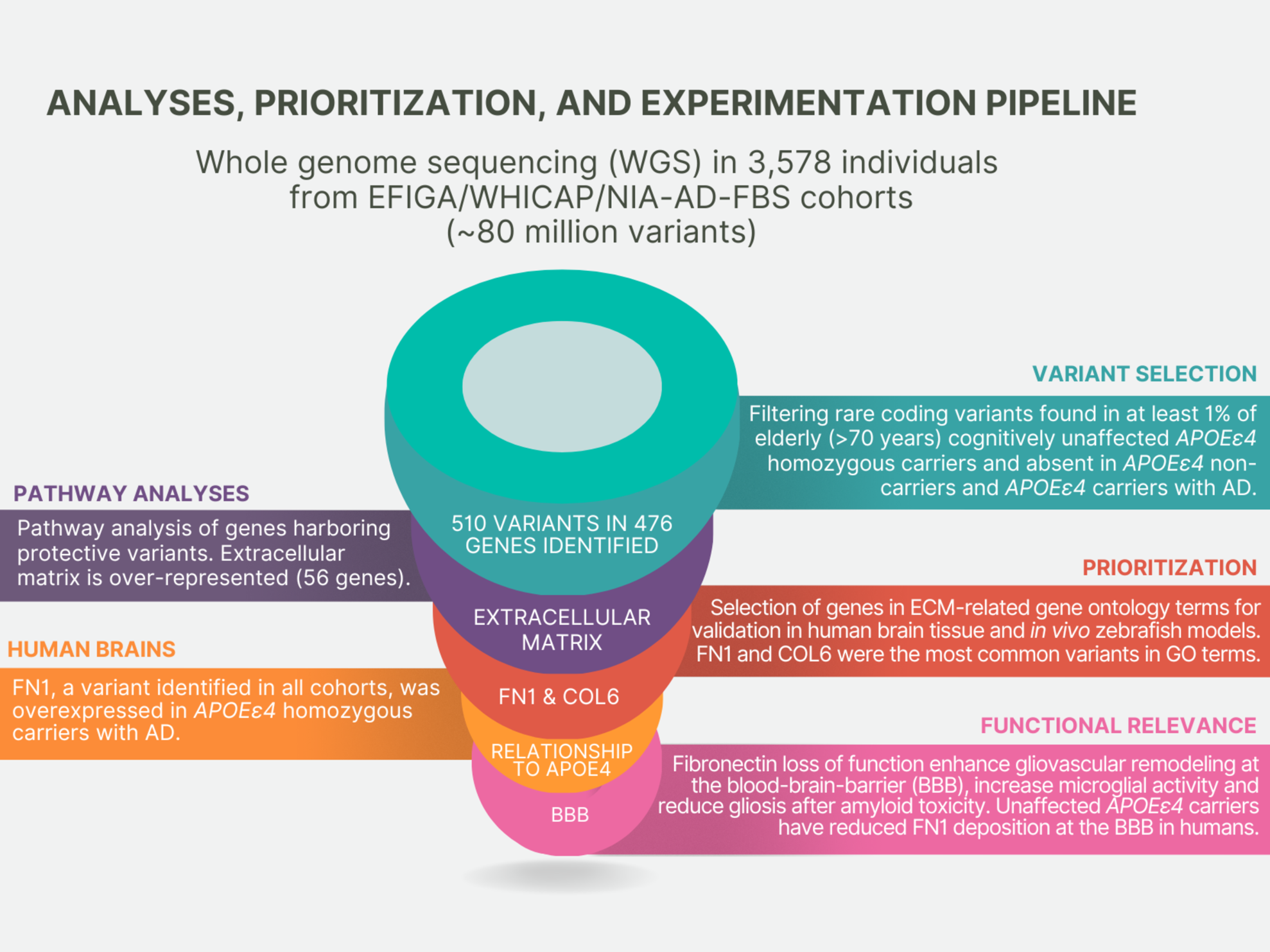
**Schematic analytical pipeline** for this study.

**Figure 3:**
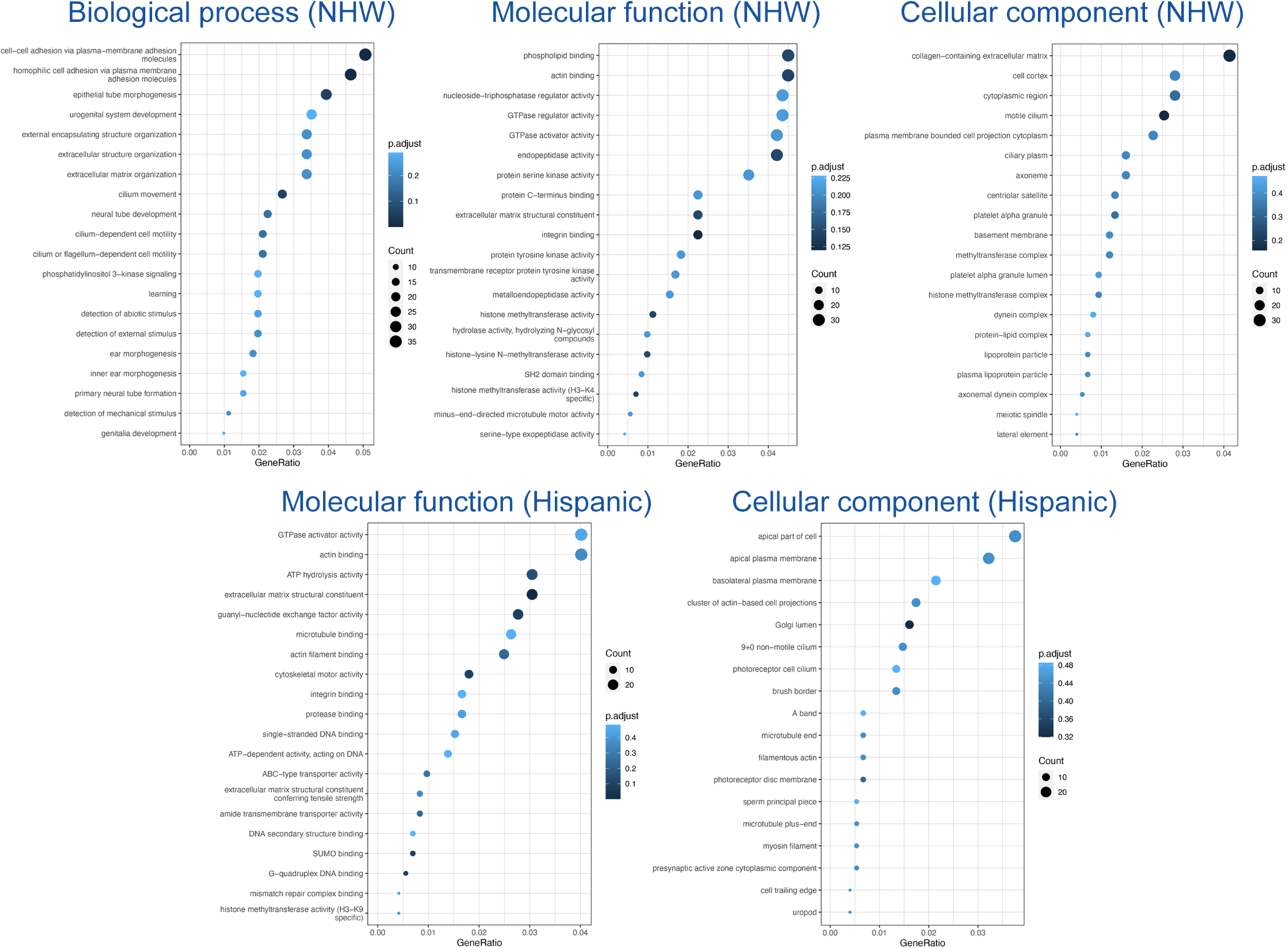
**Pathway analysis** of variants segregating in *APOEε4* carriers.

**Table 1:** Demographics of samples sequenced.

|  | Hispanics | Non-Hispanic White |
| --- | --- | --- |
| **N** | 1840 | 590 |
| AD Cases | 693 | 455 |
| AD Controls | 1147 | 135 |
| **Families with 2 or more individuals** |  |  |
| # APOE-ε4 heterozygotes | 724 | 438 |
| # APOE-ε4 homozygotes | 189 | 190 |
| # APOE-ε4 heterozygote AD cases | 442 | 265 |
| # APOE-ε4 homozygote AD Cases | 114 | 155 |
| # APOE-ε4 heterozygote healthy controls | 282 | 161 |
| # APOE-ε4 homozygote healthy controls | 75 | 30 |
| # APOE-ε4 homozygote healthy controls ≥ 70 years of age | 27 | 15 |
| # APOE-ε4 heterozygote healthy controls > 80 years of age | 75 | 45 |

**Table 2:** *FN1* minor allele frequencies.

| Cohorts | SNP | MAF in elderly* cognitively unaffected *APOEε4* homozygous | MAF in all cognitively unaffected *APOEε4* homozygous | MAF in *APOEε4* homozygous AD patients | MAF in cognitively unaffected elderly* *APOEε4* heterozygotes | MAF in all healthy *APOEε4* heterozygotes | MAF in *APOEε4* heterozygous AD patients |
| --- | --- | --- | --- | --- | --- | --- | --- |
| EFIGA/WHICAP | rs116558455 | 1.85% | 0.67% | 0.00% | 0.67% | 0.18% | 0.00% |
| NIA-FBS | rs140926439 | 3.33% | 5.17% | 0.00% | 2.22% | 1.55% | 0.96% |

\*Elderly APOE ε4 homozygous are over 70 years old and heterozygous are over 80 years old.

### Potential protective alleles against APOEε4 enrich extracellular matrix components

To determine the molecular mechanisms enriched in the protective alleles that we identified, gene ontology review as performed with term analyses for biological processes, cellular compartments, and molecular functions (**Figure 3**). We found a strong enrichment for extracellular matrix (ECM)-related processes such as cell adhesion, ECM organization, integrin binding and structural component of the ECM (**Figure 3**). This suggested that functional alterations in the ECM composition could act as a protective mechanism in *APOEε4* carriers, both heterozygotes and homozygotes without dementia. We hypothesized that *APOEε4*-related increase in ECM components could be counteracted by loss-of-function (LOF) variants in those genes, leading to protection through rescue of pathological mechanisms that those ECM components partake.

To test our hypothesis, we selected two genes from the variant lists that were common in ECM-related gene ontology classes (**Figure 3**), collagen type VI alpha 2 chain (*COL6A2*) and fibronectin 1 (*FN1*). These genes are well-known ECM components that harbored putatively protective variants in *APOEε4* cognitively unaffected carriers. Additionally, FN1 was present in both Hispanic and non-Hispanic white cohorts (**Supplementary Table 1, Supplementary Table 2**). *COL6A2* variation (rs777822883) generates a substitution of Arginine at the 862^nd^ residue to Tryptophan, while *FN1* variation (rs140926439) converts the Glycine at the 357^th^ position to Glutamic acid. Since both alterations result in change in charged residues (loss in COL6A2, gain in FN1), we hypothesized that these variations could have detrimental effect on the protein function, as charged interactions are essential for matric proteins and their stability ^11–13^. Therefore, we analyzed the AlphaFold structures of these proteins in Ensembl (www.ensembl.org) and found that both variations are potentially detrimental according to SIFT, REVEL and MetaL R predictions (**Supplementary Figure 1**). Arginine in *COL6A2* at 862^nd^ position may coordinate with Valine 859 and Glutamic acid 858 in the alpha helix structure, while Glycine at 357^th^ position in *FN1* may provide structural stability by coordinating with Glutamic acid 358 and Serine 355 (**Supplementary Figure 1**). Therefore, we categorized these variants as likely loss-of-function alleles based on loss of electrostatic interactions.

### FN1 deposition correlates with APOEε4 dosage

Based on our findings, we hypothesized that *APOEε4* dosage might correlate with deposition of COL6A2 and FN1, at the blood-brain barrier (BBB) basement membrane, one of their prominent expression locations, as FN1 is an important signaling molecule that interacts with specific integrins ^14^ expressed in various vascular niche cell types ^15^. We immunostained and analyzed the brains of 27 individuals with known *APOEε* genotypes (8 *APOEε4*/4 homozygous carriers with AD, 8 *APOEε3*/4 heterozygote carriers with AD, and 11 *APOEε4* non-carriers (*APOEε3/3*) with AD (**Supplementary Table 4**) for FN1 and CD31 (endothelial cell marker), and COL6A2 and COL4 (a vascular basement membrane marker) (**Figure 4, Supplementary Dataset 1, Supplementary Dataset 2**). We found that FN1 levels (**Figure 4A-C’**) significantly increased with *APOEε4* dosage (**Figure 4D**). Compared to *APOEε3*/3 individuals, FN1 expression increased significantly in *APOEε3/4* (8.1%, p = 3.4e-02) and in *APOEε4 homozygous* individuals (26.6%, p = 3.1e-09). Least squares linear regression and non-linear fit comparison of FN1 intensities according to the diameter of the vessels showed that compared to *APOEε*3/*3*, FN1 expression is more prominent with increasing vessel size in *APOEε*3/*4* and *APOEε4/4* individuals (adjusted R^2^: *APOEε* 3/3: 0.81, *APOEε*3/*4* 0.86, *APOEε*4/*4*: 0.89; all p values are less than 1.0xE-15 for non-zero significance of the slopes) (**Figure 4E**). Immunostainings for COL6 (**Figure 4F-H’**) showed a non-linear relationship between *APOEε4* dosage and COL6 expression. *APOEε4* heterozygotes show reduced (7.7%, p = 9.9e-03) homozygotes show increased levels of COL6 (6.7%, p = 3.4e-02) (**Figure 4I**). COL4 expression is only reduced in *APOEε4* heterozygotes (8.6%, p = 1.4e-03) but remain unchanged in homozygotes (**Figure 4I**). The changes in COL6 expression with blood vessel size was less pronounced (adjusted R^2^: *APOEε* 3/3: 0.67, *APOEε*3/*4* 0.50, *APOEε*4/*4*: 0.55; all p values are less than 1.0xE-15 for non-zero significance of the slopes) (**Figure 4J**).

**Figure 4:**
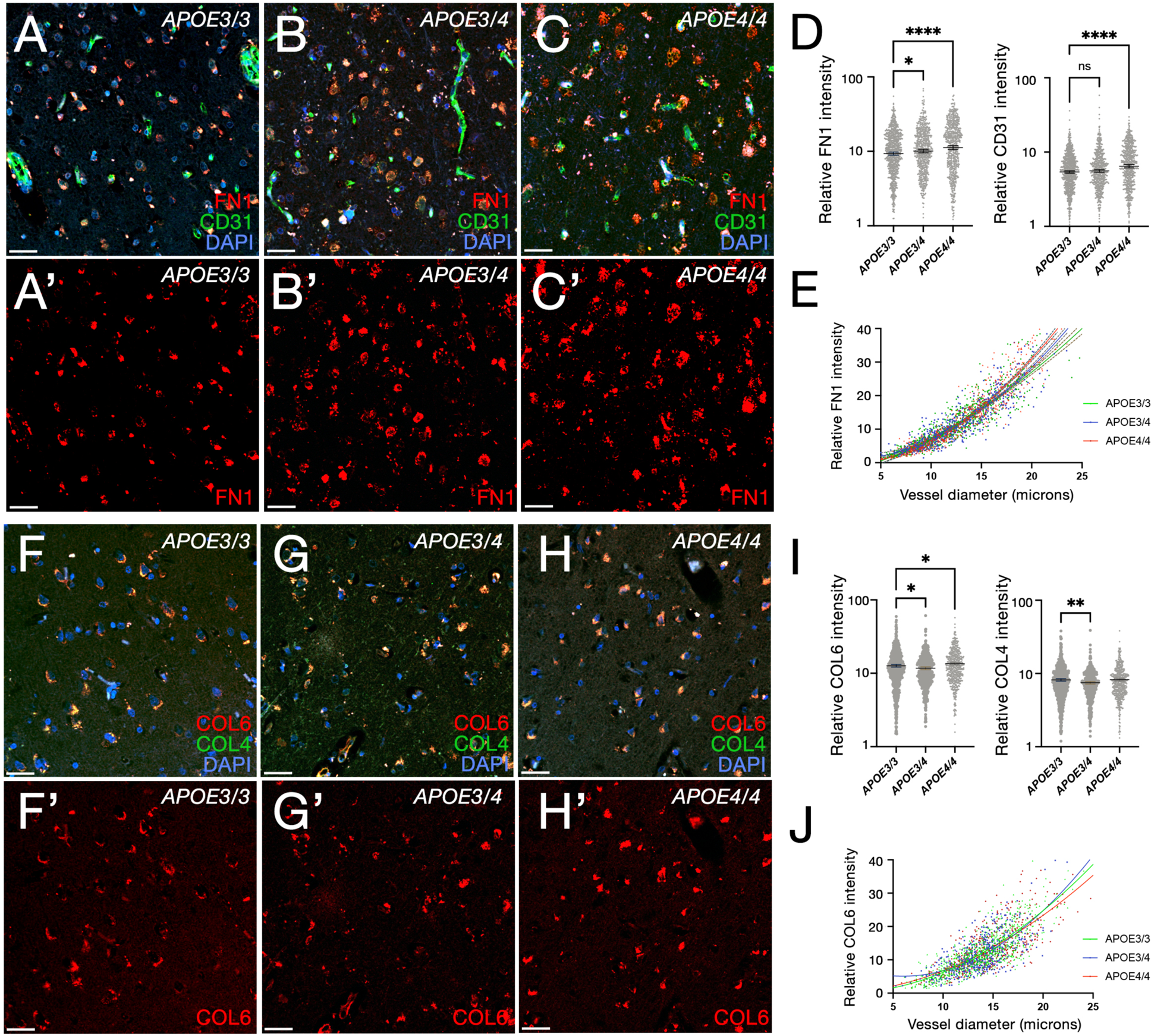
Changes in FN1 and COL6A2 according to *APOE* genotype. A-C’: Double IFS for CD31 (green) and FN1 (red) with DAPI nuclear counterstain in APOE ε3/ε3 (APOE3/3; A, A’), APOE ε3/ε4 (APOE3/4; B, B’) and APOE ε4/ε4 (APOE4/4; C, C’). D: FN1 and CD31 intensity comparisons in 2,044 blood vessels from 28 individuals. E: Regression model for FN1 intensity with respect to blood vessel diameter in three APOE genotypes. F-H’: Double IFS for COL4 (green) and COL6A2 (red) with DAPI nuclear counterstain in APOE ε3/ε3 (APOE3/3; F, F’), APOE ε3/ε4 (APOE3/4; G, G’) and APOE ε4/ε4 (APOE4/4; H, H’). I: COL4 and COL6A2 intensity comparisons in 1,816 blood vessels from 28 individuals. J: Regression model for COL6A2 intensity with respect to blood vessel diameter in three APOE genotypes.

#### FN1 deposition is different between demented and cognitively unaffected *APOEε*4/*4* carriers

Based on our findings that FN1 deposition is increased in patients with AD and *APOE* dosage correlates with FN1 levels, we hypothesized that FN1 deposition could be a downstream driver of the pathological effects of *APOEε*4 in AD. We tested this hypothesis by comparing FN1 and GFAP (marker for reactive gliosis) levels in *APOEε*3/3 (control, n = 2), *APOEε*4/4 AD (n = 2), and *APOEε*4/4 unaffected (n = 6) individuals (**Figure 5, Supplementary Table 5, Supplementary Dataset 3**). We found elevated reactive gliosis and FN1 deposition in *APOEε*4/4 carriers with AD compared to *APOEε*3/3 controls (ANOVA adjusted p=1.5E-02 for GFAP intensity, 4.1E-11 for FN1 intensity) (**Figure 5A,B,E,F,I**). *APOEε*4/4 unaffected carriers had FN1 and GFAP levels that were similar to that in controls (ANOVA adjusted p=0.5245 for GFAP intensity, p=0.8884 for FN1 intensity) (**Figure 5A,C,E,G,H**). This implies that the unaffected/resilient *APOEε*4 carriers may be protected from gliosis and FN1 deposition (**Figure 5H, I**).

**Figure 5:**
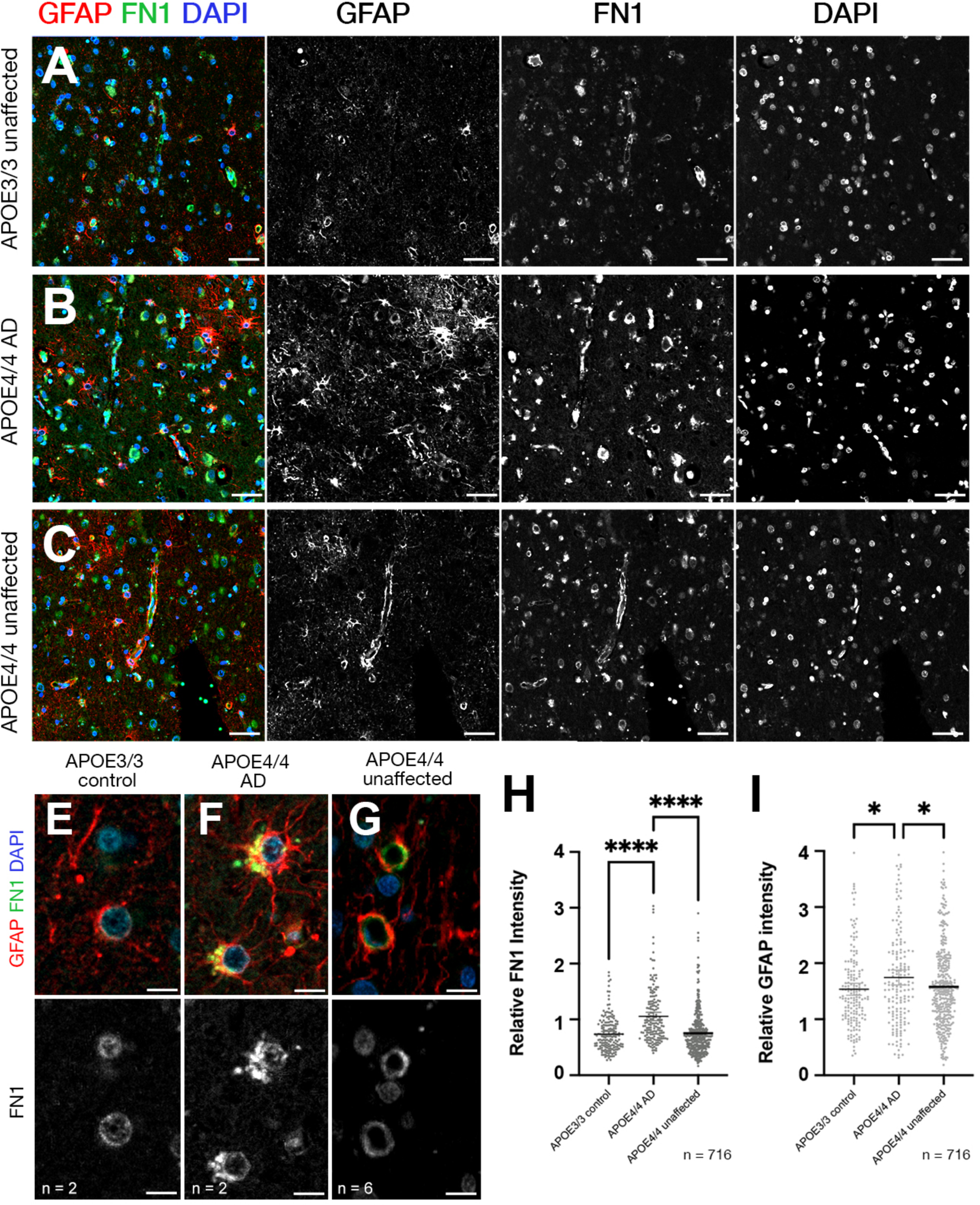
FN1 deposition and gliosis reduce to control levels in *APOEε4/4* cognitively unaffected individuals but not in *APOEε4/4* AD patients. A-C: Double IFS for FN1 (green) and GFAP (red) with DAPI nuclear counterstain in APOE ε3/3 (A), APOE ε4/4 AD (B) and APOE ε4/4 unaffected/resilient individuals (C). Black-white images are individual fluorescent channels for FN1, GFAP and DAPI. E-G: Two blood vessels in every condition are shown in high magnification together with FN1 channel alone. H: FN1 intensity comparisons. I: GFAP intensity comparisons.

#### Fibronectin loss of function zebrafish model enhance gliovascular endfeet retraction and microglial activity while reducing gliosis after amyloid toxicity

To determine whether Fibronectin activity is related to cellular responses after amyloid toxicity, we used our established amyloid toxicity model in the adult zebrafish brain ^16–21^. Zebrafish has two fibronectin 1 genes: *fn1a* and *fn1b* ^22^. Our single cell transcriptomics analyses in the zebrafish brain showed that *fn1b* but not *fn1a* is expressed in the zebrafish forebrain (**Figure 6A**). *fn1b* expression is predominantly detected in vascular smooth muscle cells and immune cells, while endothelia and astroglia expresses *fn1b* at considerably lower levels (**Figure 6B**). Amyloid toxicity results in increased *fn1b* expression in immune cells and vascular smooth muscle cells (**Figure 6B**), similar to what we observed in AD brains (**Figures 4** and **Figure 5**). To determine the effects of fibronectin function in amyloid-induced pathology, we used a *fn1b* full knockout zebrafish line (*fn1b*^−/−^), which was previously functionally validated ^23^. After treating wild type and *fn1b*^−/−^ animals with Aβ42, we performed immunohistochemical stainings for astroglia (red, GS) and tight junctions that mark vascular structures (green, ZO-1) (**Figure 6C-F, Supplementary Dataset 4**). Compared to wild type animals treated with Aβ42, *fn1b*^−/−^ animals with Aβ42 showed less co-localization of GS and ZO-1 (−16.3%, p = 5.3E-09), suggesting the gliovascular interactions are reduced with fibronectin loss of function (LOF) (**Figure 6G).** Based on our previous findings that reduced gliovascular contact upon amyloid toxicity is a protective mechanism through enhancing clearance of toxic protein aggregates and immune systems activity ^18^, our results suggest that fibronectin could be negatively regulating the amyloid beta clearance and therefore a LOF variant could be protective against disease pathology. By performing intensity measurements for astroglia with GS immunoreactivity, we observed that GS intensity reduces with *fn1b* LOF (−24.7%, p = 4.7E-03; **Figure 6H, Supplementary Dataset 5**), indicative of reduced gliotic response upon Aβ42. To determine the effect of Fibronectin on synaptic density and the number and activation state of microglia, we performed immunostainings (**Figure 6I-J**, **Supplementary Dataset 6**) and found that loss of Fibronectin leads to increased numbers of total (41.5%, p = 8.7E-04) and activated microglia (64.3%, p = 2.9E-04). We did not observe change in the synaptic density when Aβ42-treated *fn1b*^−/−^ were compared to Aβ42-treated wild type animals (**Figure 6I-K, Supplementary Dataset 7**).

**Figure 6:**
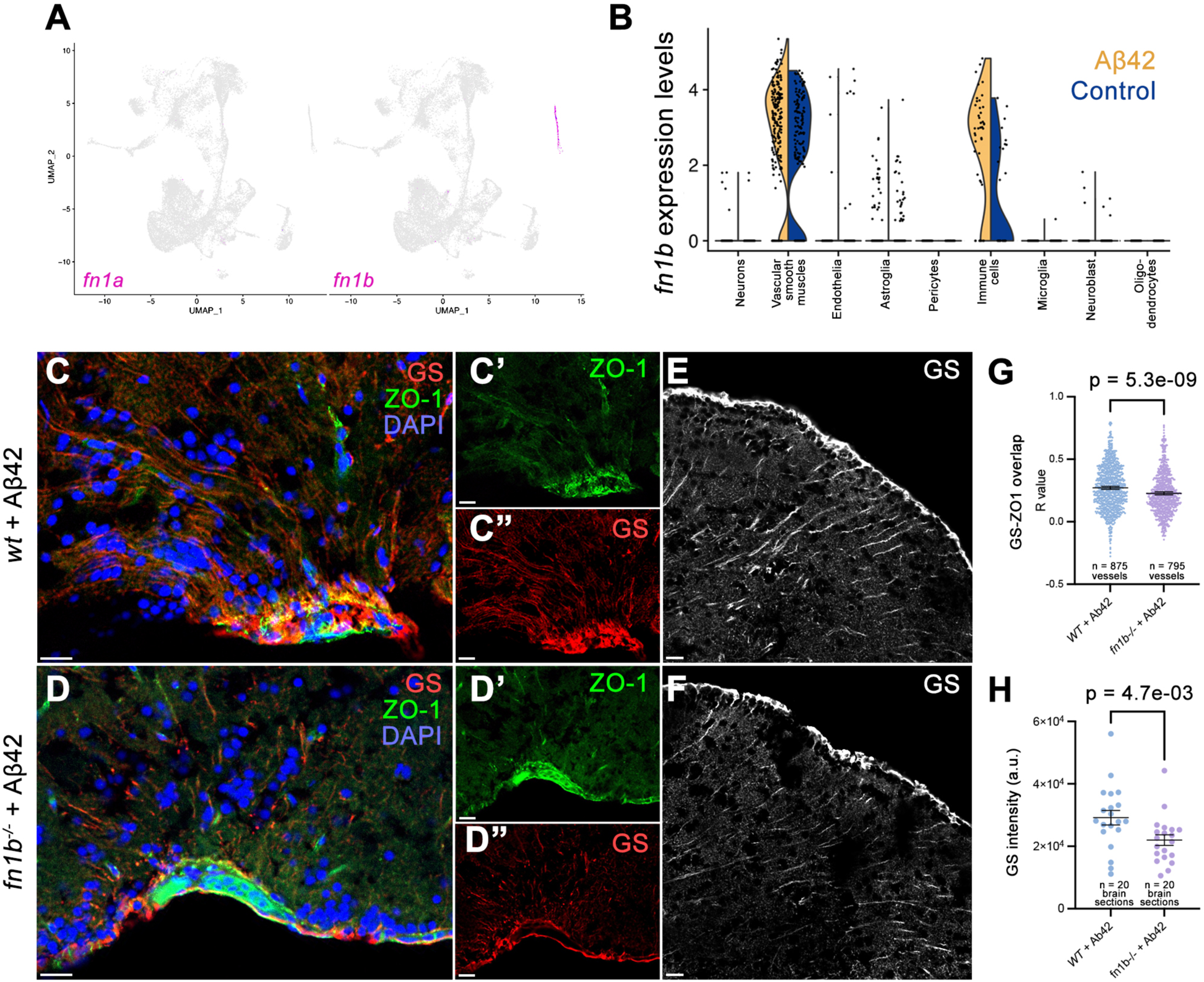
Fibronectin loss-of-function affects gliovascular interactions, gliosis, and microglial activity after amyloid toxicity in zebrafish brain. A: Feature plots for fibronectin 1a (*fn1a*) and fibronectin 1b (*fn1b*) genes in zebrafish brain. B: Violin plots in control and Aβ42-treated brains. *fn1b* is mainly expressed in vascular smooth muscle cells and immune cells and is upregulated with Aβ42. C, D: Double IF for astroglia marker glutamine synthase (GS, red) and tight junction marker (ZO-1, green) in wild type and *fn1b*^−/−^ animals. Individual fluorescent channels in C’, C’’, D’, and D’’. E, F: Individual GS channels. G: Quantification for colocalization of ZO-1 and GS. H: Comparison of intensity measurements for GS. I, J: Double IF for synaptic marker SV2 (green) and microglial marker L-Plastin (red) in wild type and *fn1b*^−/−^ animals treated with Aβ42. Individual fluorescent channels in I’, I’’, J’, and J’’. Quantifications for synaptic density, total number of microglia and activated microglia.

## Discussion

In our study, we found that two missense, potential loss-of-function (LOF) variants in *FN1* may protect against *APOEε4*-mediated AD pathology. We based our conclusions on five main observations: (1) *FN1* coding variants were present in cognitively unaffected *APOEε4* homozygous carriers but not in affected carriers with clinically diagnosed AD (**Supplementary Table 1**). (2) Deposition of FN1 at the BBB basement membrane increases with *APOEε4* dosage (**Figure 4**). (3) Unaffected/resilient homozygous *APOEε4* carriers above the age of 70 without AD have FN1 deposition levels similar to APOE*ε 3* control individuals (**Figure 5**). (4) In the zebrafish brain, knockout of *fn1b* alleviates amyloid toxicity-related pathological changes (**Figure 6**). These results suggest that the basement membrane thickening and remodeled ECM composition in the BBB may be a pathological contribution to *APOEε4*-mediated AD pathology that may be mitigated by variants in *FN1* or other ECM genes (**Figure 7**). This conclusion is supported by the presence of mutations in other BBB-related ECM components, such as *LAMA1*, *LAMA3*, and *HSPG2*, in unaffected elderly *APOEε4* carriers but not in carriers with AD (**Supplementary Table 1**). Therefore, our findings propose a new direction for potential therapeutic interventions reducing the impact of *APOEε4*-mediated risk of AD by targeting the BBB basement membrane. Thus, we propose that Fibronectin loss-of-function may be a protective mechanism for AD (**Figure 7**).

**Figure 7:**
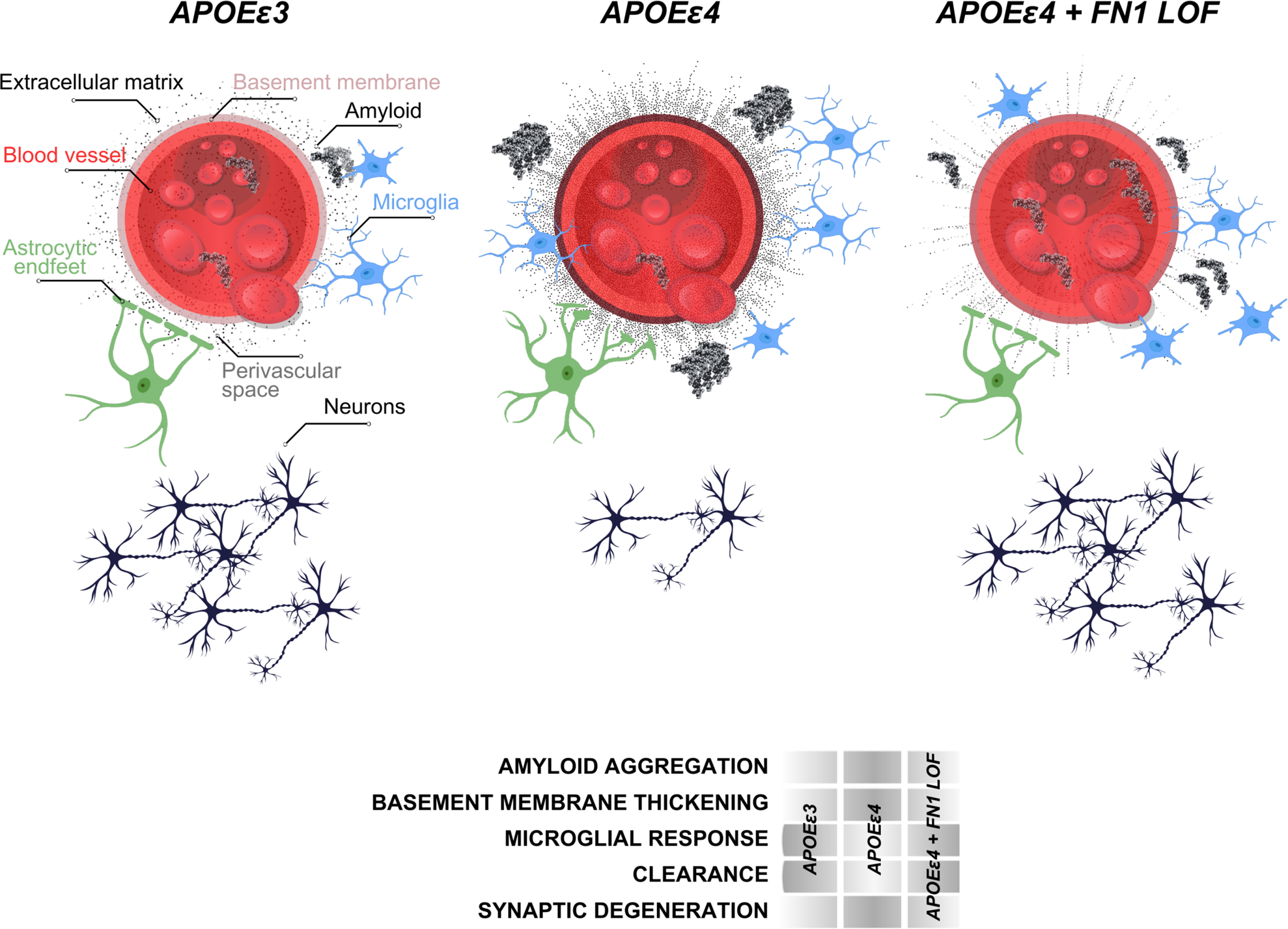
Schematic abstract for the protective effect of FN1.

*APOEε4* has been associated with increased neuroinflammation and neurodegeneration, which can accelerate the progression of AD ^24^. Our results in zebrafish *fn1b* knockout model showed that reduced Fibronectin 1 increases the gliovascular (GV) endfeet retraction and reduces the gliosis. We previously showed that the relaxed GV contact is a beneficial response to amyloid toxicity ^18^ as it helps enhance the clearance of toxic aggregates through the bloodstream. Additionally, gliosis is an immediate response in astroglia to insult, and it prevents functional restoration of neuronal activity in disease ^25–28^. Independent reports showed that astrocytic removal of *APOE* protects against vascular pathology ^29^, and gliosis is a mediator of amyloid-dependent tauopathy in late AD ^30^. We propose that the relationship of fibronectin to these processes are pathogenic, and reduced Fibronectin could be protective by allowing more efficient clearance through the bloodstream and reduced astrogliosis. The enhanced microglial activity supports this hypothesis as acute activation of microglia is a beneficial response to toxic protein aggregation ^31,32^.

Our results are consistent with the previous findings on *APOE*-dependent vascular pathologies and their relationship to AD ^8,33–37^. Endothelial fibronectin induces disintegration of endothelial integrity and leads to atherosclerotic vascular pathologies ^38–40^, supporting our findings that reduced Fibronectin 1 protects the blood-brain-barrier integrity disrupted by *APOEε4*. Our findings are coherent with the previous observations, where AD-related changes in collagen and fibronectin around the blood-brain barrier (BBB) and alterations in the BBB’s structure and function were documented ^41–43^. Additionally, the serum levels of fibronectin increase in AD patients in comparison to healthy individuals ^44^. Collagen and Fibronectin can also be early pathological markers of AD ^45^, where the increase in the deposition and crosslinking of basement membrane around cerebral blood vessels lead to a thickening of the basement membrane, potentially compromising its permeability and function ^37,46–49^. Fibronectin expression levels in brain vasculature increases in AD ^21,35,50–52^, where remodeling of the BM and replacing ECM with FN1 has been suggested to indicate hypoperfusion and atherosclerosis-prone state ^36,38,53^. Additionally, *APOEε4* might regulate the BM remodeling through inhibition of pericyte-mediated matrix proteinase expression ^9^. Pericyte degeneration, mural cell dysfunction, alterations in cerebrospinal flow dynamics are long-term consequences of vascular pathologies in aging and AD and is accelerated with *APOEε4* ^33,54–58^. Therefore, based on our findings, we propose that excess ECM deposition and BM thickening with Collagen and Fibronectin could promote blood brain barrier breakdown. Potential loss of function mutations in ECM genes are likely to render ECM components non-functional, thus protecting against AD progression. Stronger instructive interactions of collagen and fibronectin with their receptors on various BBB cell types in AD ^35,51,59,60^ support this hypothesis. Consistently, FN1 provides attachment surface for immune cells, which – when becomes chronic – damages the vascular functions, contribute to BBB breakdown and loss of synaptic integrity.

We found that despite their *APOEε4/4* status, unaffected/resilient individuals who do not develop cognitive decline have lower FN1 deposition and gliosis at the vascular basement membrane that are not different than *APOEε3/3* control individuals but significantly lower than *APOEε4/4* AD patients (**Figure 5**). This demonstrated that FN1 is a critical component of *APOEε*4-mediated development of AD, and a yet unknown protective mechanisms against the effects of *APOEε4/4* genotype suppresses FN1 deposition. We propose that FN1 is a critical downstream effector of *APOEε4*, and reduced FN1 levels, either through rare, protective genetic variations in *FN1* or through other resilience mechanisms, promotes protection against AD. An interesting future research could investigate the other rare protective variants of *APOE* such as *APOEε2* ^10,61^ and *APOEε3* Christchurch^62^ and their effects on the BBB basement membrane.

The strength of this study is the cross-species design with pathological and functional validation to show that ECM component fibronectin could be related to key pathological aspects of AD such as toxic protein clearance, blood brain barrier integrity and microglial activity. We present the first knockout zebrafish for Fibronectin 1 in relation to amyloid toxicity and identified cellular changes that relate to fibronectin activity.

Further studies could address some limitations of our study. First, the mechanism by which *APOEε4* enhances FN1 requires further investigations. Although in human and zebrafish brains, Fibronectin is upregulated, the longitudinal relationship of amyloid aggregation to FN1 activity needs to be analyzed. Additionally, our genetic studies are conducted in clinically assessed individuals, and given the rarity of the *FN1* mutation, we did not have neuropathological assessments of *APOEε4/4* individuals with this rare protective mutation. Future studies in large scale neuropathologic cohorts are necessary to demonstrate the pathological consequences of the rare *FN1* mutations. Finally, mechanistic studies of *FN1* with and without the rare mutation are necessary to demonstrate the nuanced functional consequences.

## Materials and Methods

### Ethics statement

All human samples were de-identified and the researchers cannot reach to the personal information of the donors. Institutional Review Board approval in Columbia University Irving Medical Center and Mayo Clinic was taken before the clinical data generation. Human cohorts and their characteristics are provided below. Animal experiments were carried out in accordance with the animal experimentation permits of the Institutional Animal Care and Use Committee (IACUC) at Columbia University (protocol number AC-AABN3554). Animals were maintained according to the Institutional Animal Care and Use Committee (IACUC) standards of the Institute of Comparative Medicine at the Columbia University Irving Medical Center and to the accepted guidelines^63–66^. The animal care and use program at Columbia University is accredited by the AAALAC International and maintains an Animal Welfare Assurance with the Public Health Service (PHS), Assurance number D16-00003 (A3007-01). Animal experiments were approved by the IACUC at Columbia University (protocol number AC-AABN3554). For zebrafish studies, 8-10 months old wild type AB strains or *fn1b*^−/−^ homozygous knockout fish lines of both genders were used. In every experimental set, animals from the same fish clutch were randomly distributed for each experimental condition.

### Human cohort information

#### NIA AD-Family Based Study (NIA AD-FBS)

This study recruited multiplex families across the United States. Families were included if at least one member had a diagnosis of definite or probable Alzheimer’s disease ^67,68^ with onset after age 60 and a sibling with definite, probable, or possible disease with a similar age at onset. Demographic information, diagnosis, age at onset for patients with Alzheimer’s disease, method of diagnosis, Clinical Dementia Rating Scale ^69^, and the presence of other relevant health problems was available for each individual. The age at onset for patients was the age at which the family first observed signs of impaired cognition. For unaffected family members, we used their age at the time of their latest examination without impairment. Each recruitment site used standard research criteria for the diagnosis of Alzheimer’s disease ^68^. For deceased family members who had undergone autopsy, the results were used to determine the diagnosis. For analyses, clinical Alzheimer’s disease was defined as any individual meeting NINCDS-ADRDA criteria for probable or possible Alzheimer’s disease ^68^ and definite Alzheimer’s disease when CERAD pathological criteria ^70^ was met postmortem.

#### Washington Heights/Inwood Columbia Aging Project (WHICAP)

WHICAP is a multiethnic, community-based, prospective cohort study of clinical and genetic risk factors for dementia. Three waves of individuals were recruited in 1992, 1999, and 2009 in WHICAP, all using similar study procedures ^71,72^. Briefly, participants were recruited as representative of individuals living in the communities of northern Manhattan and were 65 years and older. At the study entry, each person underwent a structured interview of general health and function, followed by a comprehensive assessment including medical, neurological, and psychiatric histories, and standardized physical, neurological, and neuropsychological examinations. Individuals were followed every 18-24 months, repeating examinations that were similar to baseline. All diagnoses were made in a diagnostic consensus conferences attended by a panel consisting of at least one neurologist and one neuropsychologist with expertise in dementia diagnosis, using results from the neuropsychological battery and evidence of impairment in social or occupational function. All-cause dementia which was determined based on *Diagnostic and Statistical Manual of Mental Disorders, 4th Edition criteria* ^73^. Furthermore, we used the criteria from the National Institute of Neurological and Communicative Disorders and Stroke–Alzheimer Disease and Related Disorders Association to diagnose probable or possible AD^68^.

#### Estudio Familiar de Influencia Genetica en Alzheimer (EFIGA)

We used families from a different ethnic group to identify protective alleles in *APOEε4* healthy individuals. This cohort comprises of participants from a group of families from the Dominican Republic, Puerto Rico, and New York. Recruitment, study design, adjudication, and clinical assessment of this cohort was previously described ^74^ as were details of genome-wide SNP data, quality control and imputation procedures of the GWAS data ^75,76^. Participants were followed every two years and evaluated using a neuropsychological battery^77^, a structured medical and neurological examination and an assessment of depression ^78,79^. The Clinical Dementia Rating Scale (CDR) ^80,81^ and functional status were done and the clinical diagnosis of Alzheimer’s disease was based on the NINCDS-ADRDA criteria ^82,83^.

### Whole genome sequencing and quality control

The demographics of the individuals selected for sequencing is shown in **Table 1**. WGS was performed at the New York Genome Center (NYGC) using one microgram of DNA, an Illumina PCR-free library protocol, and sequencing on the Illumina HiSeq platform. We harmonized the WGS the EFIGA families (n=307), and jointly called variants to create a uniform, analysis set. Genomes were sequenced to a mean coverage of 30x. Sequence data analysis was performed using the NYGC automated analysis pipeline which matches the CCDG and TOPMed recommended best practices ^84^. Briefly, sequencing reads were aligned to the human reference, hs38DH, using BWA-MEM v0.7.15. Variant calling was performed using the GATK best-practices. Variant filtration was performed using Variant Quality Score Recalibration (VQSR at tranche 99.6%) which identified annotation profiles of variants that were likely to be real and assigns a score (VQSLOD) to each variant.

### Identification of variants segregating in healthy *APOEε4* individuals

First, we filtered high quality rare (MAF<0.01 in GnomAD) variants with genotype quality (GQ)≥20 and depth (DP)≥10. We then excluded any variant observed in *APOE* ε4 non-carriers. Within variants that segregated in *APOE*ε4 carriers, we prioritized those that were observed in at least one *APOEε4* homozygous healthy elderly (≥70 years) and had additional support in healthy elderly (≥80 years) heterozygous carriers. We further prioritized variants that were absent in AD patients carrying an *APOEε4* allele. A simplified pipeline is provided in **Figure 2**.

### Genotyping, amyloid administration, tissue preparation

A previously generated *fn1b* knockout line using CRISPR-Cas9 gene editing ^23^ was used in homozygous form. The full deletion was genotyped as described ^23^. Amyloid-β42 was administered to the adult zebrafish brain through cerebroventricular microinjection into the cerebral ventricle ^20^. Euthanasia and tissue preparation were performed as per institutional ethic committee approval and international guidelines ^20,65^. 12-µm thick cryo-sections were prepared from these brain samples using a cryostat and collected onto glass slides which were then stored at −20°C.

### Immunohistochemistry

Post-mortem human brain sections from BA9 prefrontal cortex were obtained from New York Brain Bank at Columbia University and Mayo Clinic Jacksonville as paraffin-embedded blanks and with neuropathology assessments (**Supplementary Tables 4-5**). Immunohistochemistry (IHC) was performed as described ^18,85^. As primary antibodies FN1 (Proteintech, catalog number 66042-1-Ig, 1:250), CD31 (Abcam, catalog number ab134168, 1:250), COL6A2 (Thermofisher, catalog number PA5-65085, 1:200), COL4 (Thermofisher, 14-9871-82, 1:100), and GFAP (Thermofisher, catalog number OPA1-06100), and as secondary antibodies goat anti-mouse Alexa Fluor 448 (Thermofisher, catalog number A-21131, 1:500) and goat anti-rabbit Alexa Fluor 555 (Thermofisher, catalog number A-21137, 1:500) were used. In short, for deparaffinization and hydration, xylene and alcohol were used. Antigen retrieval was performed with citrate buffer (pH:6.0) or Antigen Retriever EDTA buffer (pH:8.5) in a pressure cooker or microwave for 18-25 minutes. Sections were blocked in 10% normal goat serum for 1 hour at room temperature and were incubated with primary antibody combinations (FN1-CD31, COL6A2-COL4 or FN1-GFAP) overnight at 4°C in a humidified chamber. Each secondary antibody to respective primaries were applied for 2 hours at room temperature. Slides were covered by mounting medium with nuclear counterstain DAPI (Thermofisher, catalog number D1306, 5 ng/ml). Immunohistochemistry for zebrafish was performed as described^20^. In short, the slides were dried at room temperature for 30 minutes and washed with PBS with 0.03% Triton X-100 (PBSTx). Primary antibodies combinations (ZO-1 + GS and SV2A + L-Plastin) were applied overnight at 4°C. Next day, after 3 times with PBSTx appropriate secondary antibodies were applied for 2 hours at room temperature. The slides were then washed several times before mounting using 70% glycerol in PBS. The following antibodies were used: mouse anti-ZO-1 (1:500, Thermofisher Cat. No. 33-9100), rabbit anti-Glutamine synthetase (GS) (1:500, Abcam Cat. No. ab176562), mouse anti-SV2A (1:500, DSHB Cat. No. SV2), and rabbit anti-L-Plastin (1:3000, gift from Michael Redd), secondary antibodies goat anti-mouse Alexa Fluor 448 (Thermofisher, catalog number A-21131, 1:500) and goat anti-rabbit Alexa Fluor 555 (Thermofisher, catalog number A-21137, 1:500). For antigen retrieval of ZO-1 and SV2, slides were heated in 10mM Sodium acetate at 85°C for 15 minutes before primary antibody incubation.

### Image acquisition, quantification, statistical analyses

Five random illumination field images per patient from the immunostained slides were acquired using Zeiss LSM800 confocal microscope equipped with ZEN software (version blue edition, v3.2, Carl Zeiss, Jena, Germany). Based on vascular markers, coronally sectioned blood vessels were delineated with the selection tool of ZEN software. Fluorescence intensity measures, diameter and area was calculated. Acquisitions were performed in blinded fashion (sample IDs, neuropathology details and genotypes were revealed after the acquisition) and no vessels were specifically left out unless their diameters were larger than 50 μm. GraphPad Prism software version 9.2.0. was used for the statistical analyses. For multiple comparisons, one-way Brown-Forsythe and Welch ANOVA test with two-stage linear step-up procedure of Benjamini, Krieger and Yekutieli comparison with individual calculation of variances was used. For non-Gaussian distributions, non-parametric Kruskal-Wallis test with Dunn’s multiple comparison test was performed. For correlation of vessel diameter to fluorescent intensity, simple linear regression model and second order polynomial robust regression with no weighting was used. Significance is indicated by ∗ (p < 0.0332), ∗∗ (p < 0.0021), ∗∗∗ (p < 0.002), **** (p<0.0001). No asterisks indicate non significance. No sample set was excluded from the analyses unless the histological sections were damaged severely during the acquisition of the sections (constitutes less than 3% of all sections analyzed).

For zebrafish studies, the effect sizes for animal groups were calculated using G-Power, and the sample size was estimated with n-Query. 4 zebrafish from both sexes were used per group. For quantification of SV2-positive synapses, 3D object counter module of ImageJ software was used with a same standard cut-off threshold for every image. For quantification of activated/resting L-Plastin-positive microglial cells, two different microglial states were classified based on their cellular morphology: slender and branched as resting microglia; round and regular as active microglia. 6 images each from telencephalon sections were analyzed per animal. For colocalization studies, vascular fields were determined using ZO-1 staining on sections (20 for every group), and colocalization with glial endfeet labelled with GS stainings was performed by using ImageJ software (v.2.1.0/1.53c) with its Colocalization Test. Data acquisition was randomized with Fay (x,y,z translation) to acquire in total 1,670 data points from two experimental groups. R(and) correlation values from wild type and *fn1b*^−/−^ animals were compared using GraphPad Prism (v.9.2.0). Intensity values for individual fluorescent channels were obtained with modal gray value and integrated density measurements using ImageJ. Comparison of 40 sections from two experimental groups was performed. Unpaired non-parametric Kolmogorov-Smirnov t-test was performed for testing the statistical significance for all analyses.

### In silico structure prediction

Protein structures, interspecies similarities and the deleterious effects of mutations were analyzed by SWISS-MODEL protein structure homology-modelling server through Expasy web server (https://swissmodel.expasy.org). SWISS-MODEL repository entry for respective proteins were retrieved and compared to desired protein orthologs using the superposition function. Deleterious mutation prediction was performed using Ensembl-integrated AlphaFold prediction model with SIFT, MetaLR and REVEL modules of prediction of deleteriousness.

### Amyloid toxicity and single cell sequencing

Amyloid toxicity was induced as described^18,20^ in the adult telencephalon, the brains were dissected and single cell suspensions were generated as previously described^86,87^. Chromium Single Cell 3’ Gel Bead and Library Kit v3.1 (10X Genomics, 120237) was used to generate single cell cDNA libraries. Generated libraries were sequenced via Illumina NovaSeq 6000 as described^20,86–89^. The cell clusters were identified using a resolution of 1. In total, 34 clusters were identified. The main cell types were identified by using *s100b* and *gfap* for astroglia; *sv2a, nrgna, grin1a, grin1b* for neurons; *pdgfrb* and *kcne4* for pericytes; *cd74a* and *apoc1* for microglia; *mbpa* and *mpz* for oligodendrocytes; *myh11a* and *tagln2* for vascular smooth muscle cells, *kdrl* for endothelial cells^10,15^. The zebrafish gliovascular single cell dataset can be accessed at NCBI’s Gene Expression Omnibus (GEO) with the accession number GSE225721.

## Author contributions

R.M., C.K. and B.N.V. conceived and designed this study. T.I.G and B.N.V. generated the variations list. D.R-D., R.L., M.M., D.R., P.R., R.M. provided the AD cohort information. N.E-T., D.W.D., D.F., and A.F.T. provided the human brain samples. D.J. and S.H. provided the *fn1b* knockout zebrafish. P.B. and C.K. performed the human postmortem tissue immunohistochemistry, image acquisition, quantification, and statistical analyses. E.Y., P.B., C.K. performed single cell sequencing, data analyses and interpretation. P.B., H.T., C.K performed the amyloid toxicity assessment and relevant quantifications in zebrafish brains. R.M., B.N.V, and C.K. interpreted the results. C.K., B.N.V, and R.M. wrote the manuscript. All authors read, edited, and approved the final version.

## Acknowledgements

We thank the Carol and Gene Ludwig Family Foundation and the Agouron Institute for providing grant support for this work. We would like to thank Taub Institute for Research on Alzheimer’s Disease and the Aging Brain Imaging Platform at Columbia University, Molecular Pathology (MPSR) and Flow Cytometry Core Facility (CCTI, supported in part by the Office of the Director, National Institutes of Health under awards S10OD020056) platforms of the Columbia University Herbert Irving Comprehensive Cancer Center for procedural support, and New York Brain Bank for post-mortem human brain sections. We thank the contributors, who collected samples used in this study. We thank the patients and families for their participation, without whom these studies would not have been possible.

This work was supported by National Institute on Aging R01 AG067501 (Genetic Epidemiology and Multi-Omics Analyses in Familial and Sporadic Alzheimer’s Disease Among Secular Caribbean Hispanics and Religious Order) (R.M, B.N.V, C.K.) and National Institute on Aging RF1 AG066107 Epidemiological Integration of Genetic Variants and Metabolomics Profiles in Washington Heights Columbia Aging Project (R.M., B.N.V, C.K), NIGMS grant R35GM148348 (S.A.H), Schaefer Research Scholars Award (C.K.), Taub Institute Grants for Emerging Research (TIGER) (C.K.) and The Thompson Family Foundation Program for Accelerated Medicine Exploration in Alzheimer’s Disease and Related Disorders of the Nervous System (TAME-AD) (C.K.). N.E-T is supported by NIH/NIA grants R01AG061796, U19AG074879, U01AG046139 and is recipient of the Alzheimer’s Association Zenith Award. We thank the Carol and Gene Ludwig Foundation to their grant support to B.N.V. to conduct this study.

The National Institute on Aging-AD Family-based study (NIA AD-FBS; https://www.neurology.columbia.edu/research/research-centers-and-programs/national-institute-aging-alzheimers-disease-family-based-study-nia-ad-fbs) collected the samples used in this study and is supported by National Institute on Aging (NIA) grants U24AG026395, U24AG021886, R01AG041797, and U24AG056270. Additional families were contributed to the NIA-AD FBS through NIH grants: R01AG028786, R01AG027944, RO1AG027944, RF1AG054074, U01AG052410. The NIA-AD FBS began in 2003 with the goal of recruiting large, multiply-affected families with late-onset Alzheimer’s disease (AD) for genetic research. The study created a resource of well-characterized families with late-onset AD. The initial phases of the Alzheimer’s Disease Sequencing Project (ADSP) included genotyping of hundreds of participants from NIA-AD FBS. The ADSP Follow-Up Study heavily engages resources provided by the NIA-AD FBS and depends upon the longitudinal follow-up of families, and the collection of additional families, in particular those from diverse populations. Samples include biological materials for genome wide association studies (GWAS) and whole genome sequencing (WGS), peripheral blood mononuclear cells (PBMC) for stem cell modeling, plasma for studies of metabolomics, proteomics, and biomarker research, and brain autopsy materials for bulk RNA sequencing.

WHICAP Data collection and sharing for this project was supported by the National Institutes on Aging (NIA) of the National Institutes of Health (NIH): R01AG072474, AG066107, AG059013.

Estudio Familiar de Influencia Genetica en Alzheimer (EFIGA) is a study of sporadic and familial Alzheimer’s Disease among Caribbean Hispanics recruited from clinics in the Dominican Republic and New York (R01 AG067501). The goal of this study is to identify genetic variants that increase late onset Alzheimer disease risk in this ethnic group. This study was initiated in 1998 and recruited individuals and their families in New York as well as from clinics in the Dominican Republic. Recruitment for the EFIGA began in 1998, to study the genetic architecture of AD in the Caribbean Hispanic population. Patients with familial AD were recruited and if a sibling of the proband had dementia, all other living siblings and available relatives underwent evaluation. Cases were defined as any individual meeting NINCDS-ADRDA criteria for probable or possible AD. EFIGA study is supported by NIA grants R56AG063908, R01AG067501 and RF1AG015473. We acknowledge the services of CEDIMAT for collaborating with sample collection and processing in the EFIGA cohort.

The content of this publication is solely the responsibility of the authors and does not necessarily represent the official views of the National Institutes of Health.

---

**Supplementary Figure 1:**
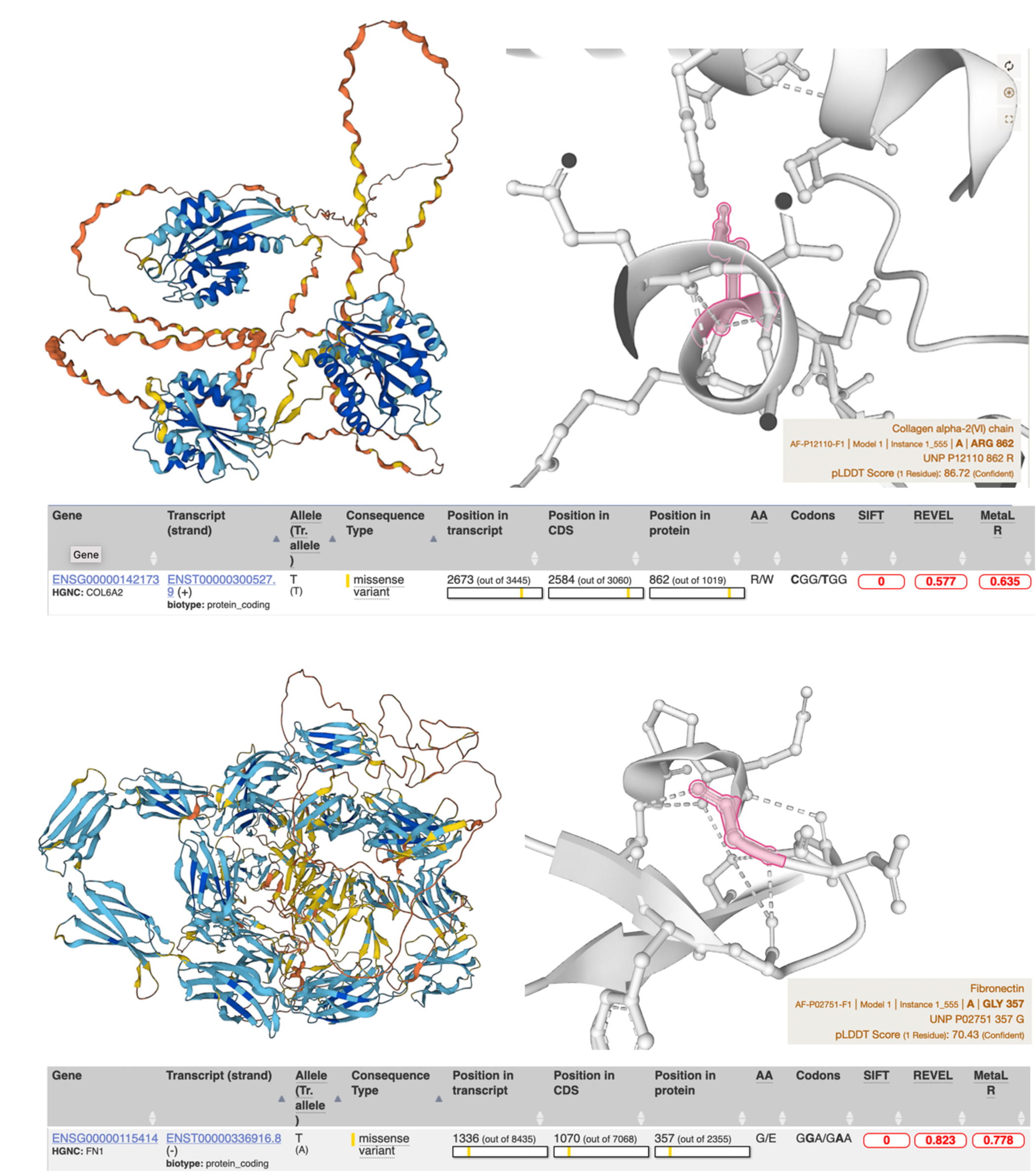
Structure and deleteriousness prediction for FN1 and COL6A2.

